# Human hippocampal ripples loosen medial prefrontal constraints to foster creative thinking

**DOI:** 10.1101/2025.07.22.666046

**Authors:** Li He, Xiongfei Wang, Jinbo Zhang, Zhibing Xiao, Xiangyu Hu, Yunzhe Liu

## Abstract

Creative thinking often requires breaking away from dominant mental schemas to generate novel and useful ideas. Yet how the human brain relaxes schematic constraints to enable creativity remains unclear. Using human intracranial EEG recordings, we found that moments of creative idea generation were accompanied by increased hippocampal ripple events, a concurrent reduction in ripple-triggered high-frequency activity in the medial prefrontal cortex (mPFC), and greater divergence of ripple-triggered mPFC activity patterns from the typical, schema-like task state. These effects followed a dorsal–ventral gradient within the mPFC and were most pronounced in the ventromedial PFC (vmPFC), a region heavily involved in schematic representation. Together, our findings suggest that hippocampal ripples can transiently disengage medial prefrontal schema constraints, thereby promoting more flexible and creative thinking.

## Introduction

We tend to default to familiar schemas, mental models of how things typically function (Bartlett, 1932; Rumelhart, 1980). While schemas are helpful for routine tasks, they can also constrain creativity (Chrysikou & Weisberg, 2005; Duncker, 1945; Knoblich et al., 1999). For example, the functional fixation on viewing a watermelon solely as food (e.g., for eating or juicing) can limit our ability to imagine unconventional uses, such as a flowerpot or helmet. The hippocampus and the default mode network (DMN), particularly the medial prefrontal cortex (mPFC), are thought to play a key role in enabling the shift from the familiar to the inventive (Bartoli et al., 2024; Becker & Cabeza, 2025; Bendetowicz et al., 2018; Duff et al., 2013; Luchini et al., 2025; Luo & Niki, 2003; Ren et al., 2020; Shofty et al., 2022). Strong hippocampus–mPFC coupling facilitates schema-based reasoning and memory retrieval (Audrain & McAndrews, 2022; den Bakker et al., 2023; Sekeres et al., 2024; Xiao et al., 2025), but may also reinforce rigid responses (Guise & Shapiro, 2017; Zeithamova et al., 2012). By contrast, weakening this coupling can “reset” entrenched habitual patterns and restore flexibility (Park et al., 2021). However, how the brain dynamically modulates hippocampus– mPFC interactions to support creative thought remains unknown.

A promising neural mechanism for such modulation is the hippocampal ripple, a brief burst of high-frequency oscillatory activity in the hippocampus (Buzsáki, 2015). Hippocampal ripples have been extensively linked to memory consolidation (Bush et al., 2022; Zhang et al., 2018), retrieval (Norman et al., 2019), inference (Xiao et al., 2025), and spontaneous thought (Iwata et al., 2024). They are thought to support these diverse cognitive functions in part by coordinating the replay of experience-related neural sequences (Buzsáki, 2015; Ólafsdóttir et al., 2018). Such replay sequences can represent exact replays of prior experiences, but can also take on a more constructive role – for example, by simulating novel scenarios in which disparate memories and knowledge are flexibly recombined (Y. Liu et al., 2019; Schwartenbeck et al., 2023). This “generative power” of hippocampal ripples has been proposed as a potential neural substrate for creative thought (Aru et al., 2023; Lewis et al., 2018).

Hippocampal ripples act as a transient gating signal, delivering brief bursts of excitatory input to widespread neocortex, momentarily reshaping activity patterns in DMN hubs (Buzsáki, 2015; Nitzan et al., 2022; Norman et al., 2021; Xiao et al., 2025). Ripple-triggered reconfiguration may disengage prefrontal constraints, enabling access to remote or unusual associations (Kucyi et al., 2023; Mildner & Tamir, 2019). Consistent with this view, disrupting sustained hippocampal–mPFC coupling in animals facilitates the learning of new rules by breaking habitual patterns (Park et al., 2021). Similarly, in humans, ripple events may promote a more flexible exploration of conceptual space and spur the emergence of original ideas (Aru et al., 2023). We hypothesised that hippocampal ripples could intermittently relieve prefrontal “schematic control,” fostering more flexible and creative thought.

## Results

### Hippocampal ripples are engaged during creative thinking and predict everyday creativity

To test this hypothesis, we recorded intracranial EEG (iEEG) simultaneously from the hippocampus (247 electrode contacts) and widespread cortical regions – including 815 contacts in the DMN – in 40 neurosurgical patients (**table S1**). Participants performed a classic creative thinking task, the Alternative Uses Task (AUT), where they had to invent unconventional uses for common objects (Guilford, 1967; Torrance, 1962), and a control task called the Object Characteristics Task (OCT), which required listing each object’s typical features (Fink et al., 2009). Participants remained silent during each response’s think period and verbalised each idea aloud immediately when it came to mind. The number of responses did not differ significantly between the two tasks (β = - 0.263 ± 0.140, *P* = 0.062; **Fig. 1A**).

**Fig. 1.**
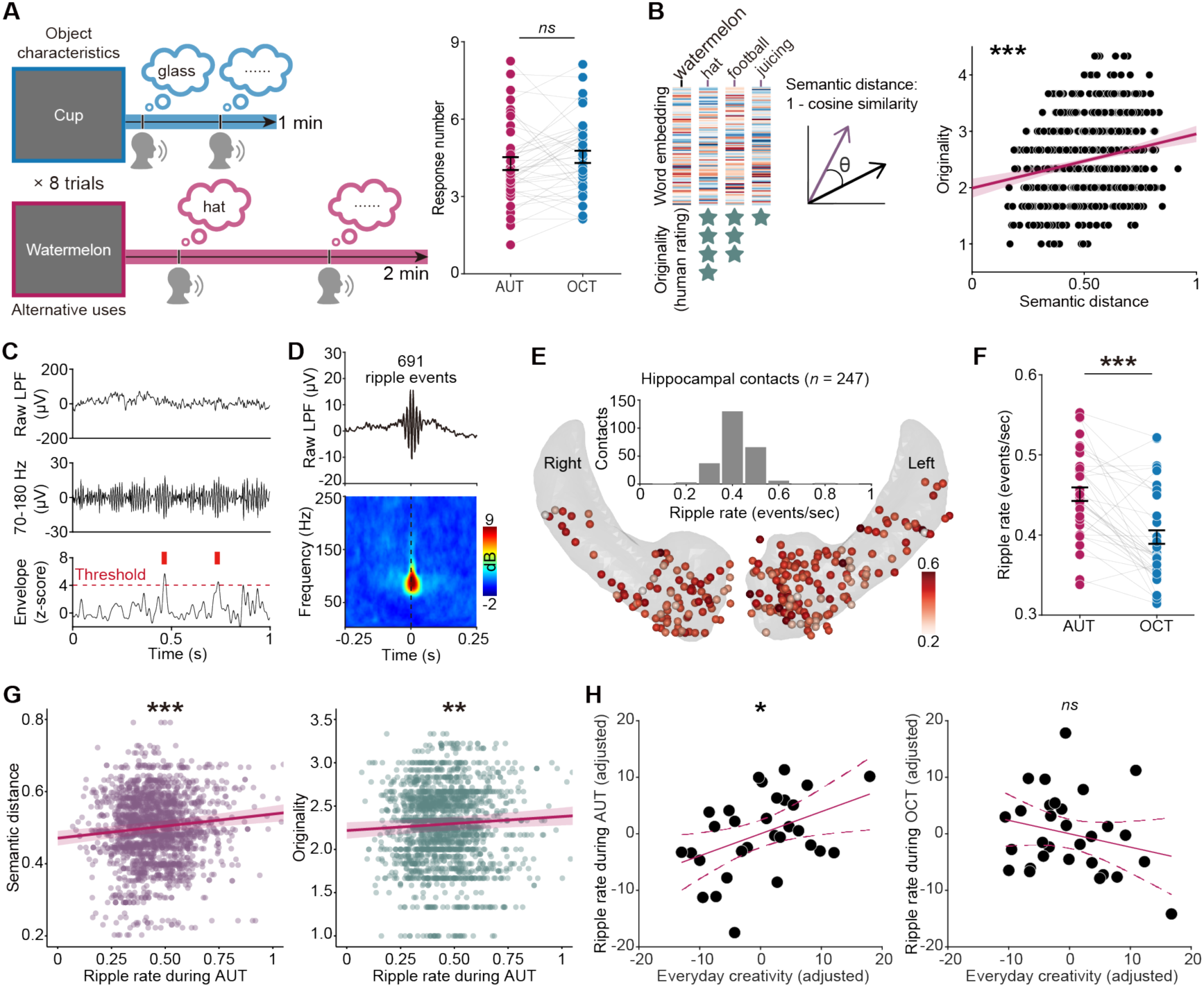
Hippocampal ripples are engaged during creative thinking. (**A**) Task schematic. In the control OCT, subjects generated typical features of a common object (e.g., “made of glass” for cup), whereas in the AUT they generated novel uses for the object (e.g., using a watermelon as a “hat”). The number of responses was similar between the two tasks. Data show mean ± s.e.m, estimated using an LME model. Dots represent individual subjects for illustration purposes. (**B**) Semantic distance between AUT responses and the target object (defined as 1 minus cosine similarity of their word embeddings) was positively correlated with human-rated originality (814 unique responses across 8 objects). (**C**) Example of hippocampal ripple detection. Red tick marks indicate ripple events detected on a hippocampal contact (see Methods for details). (**D**) Mean peri-ripple field potential from a hippocampal contact (averaged across all its ripple events) and corresponding wavelet spectrogram centred at the ripple peak. (**E**) Location of hippocampal recording contacts in all 40 subjects, with each contact’s marker coloured according to its overall ripple rate during the task. Inset: histogram of ripple rates across hippocampal contacts. (**F**) Hippocampal ripple rate was higher during the AUT than the OCT. Data show mean ± s.e.m, estimated using an LME model. Dots represent individual subjects for illustration purposes. (**G**) Across trials, higher hippocampal ripple rates were associated with greater semantic distance and higher originality of AUT responses. Shaded bands indicate the 95% confidence interval of the regression line, estimated using an LME model. (**H**) Hippocampal ripple rate during the AUT positively correlated with everyday creativity (questionnaire score), controlling for ripple rate during the OCT and Raven’s IQ score. No such association was observed during the OCT. Dashed bands indicate the 95% confidence interval of the fitted regression line. Dots represent individual participants. \**P* < 0.05; \*\**P* < 0.01; \*\*\**P* < 0.001.

All responses were recorded, transcribed, and time-stamped using the audio signal for alignment with neural data. To quantify response creativity, three independent raters scored each AUT response for originality on a 5-point scale (with good inter-rater reliability, intraclass correlation = 0.68). We also computed the semantic distance of each AUT response from its object prompt (defined as 1 minus the cosine similarity between their word embeddings in a 100-dimensional Chinese corpus model (Y. Song et al., 2018)). Only meaningful verbal responses were retained for analysis (see examples in **table S2**); any off-topic responses (as manually identified by raters) were excluded. For the remaining responses, semantic distance was strongly correlated with human-rated originality (Pearson *r* = 0.215, *P* < 10^-10^; **Fig. 1B**), confirming it as a useful objective index of creative output (Heinen & Johnson, 2018).

Hippocampal ripple events were identified by their characteristic high-frequency oscillatory waveform (**Fig. 1, C and D**; see Methods for details). Critically, hippocampal ripples were significantly more frequent during creative idea generation than during the control task. Ripple rates in the AUT were higher than in the OCT (linear mixed-effects, LME, β = 0.053 ± 0.004, *P* < 10^-41^; **Fig. 1F**). Furthermore, trials with more ripples tended to produce more creative responses: within the AUT, higher ripple rates were associated with larger semantic distances (β = 0.067 ± 0.015, *P* < 10^-6^) and with higher originality ratings of the responses (β = 0.162 ± 0.061, *P* = 0.009; **Fig. 1G**). Together, these findings provide the direct evidence that hippocampal ripples are engaged during creative thinking.

We also tested whether an individual’s average ripple rate during the creative-thinking task could predict real-world creative tendencies. Participants completed a validated inventory measuring the frequency of everyday creative activities across various domains (e.g., literature, music, cooking), providing a behavioural index of creativity in daily life (Benedek et al., 2020). We also administered Raven’s Standard Progressive Matrices (Raven, 2000) to control for general fluid intelligence. Notably, hippocampal ripple rate during the AUT was positively associated with everyday creative activity (*r* = 0.419, *P* = 0.026; **Fig. 1H**), after accounting for ripple rate during the OCT and Raven’s score. In contrast, hippocampal ripple rate during the OCT was not associated with everyday creativity (*r* = −0.237, *P* = 0.224). The correlation between ripple rate during the AUT and everyday creativity was significantly stronger than that during the OCT (*z* = 5.836, *P* < 0.001). As a control, hippocampal ripple rate during neither the AUT nor the OCT was significantly related to Raven’s score (AUT: *r* = 0.004, *P* = 0.984; OCT: *r* = −0.020, *P* = 0.918). These findings suggest that hippocampal ripples during creative thinking also reflect individual differences in real-world creative activity, beyond general intelligence or ripple occurrence in non-creative contexts.

### Ripple-aligned high-frequency activity in the mPFC supports creative thinking

In the neocortex, regions of the DMN are consistently recruited when people engage in imaginative and creative thought (Bartoli et al., 2024; Beaty et al., 2016; Luchini et al., 2025; Shofty et al., 2022). We therefore examined cortical high-frequency broadband activity (HFB, 60–120 Hz power, a proxy for local neuronal firing (Parvizi & Kastner, 2018)), time-locked to hippocampal ripples (±250 ms centred on the hippocampal ripple peak) (Norman et al., 2021), focusing on the DMN and its subregions.

We used a linear regression-based deconvolution approach (Ehinger & Dimigen, 2019; Norman et al., 2021), to disentangle ripple-locked cortical activity from other possible overlapping events (e.g., the onset or offset of a verbal response), since such events could occur in close temporal proximity (e.g., a hippocampal ripple might occur shortly before or after a vocalised idea). Continuous HFB power time series in each cortical contact were modelled with time-expanded predictors aligned to the timing of all relevant ripple events, enabling the estimation of ripple-specific HFB responses with shared variance appropriately partitioned across overlapping events (see Methods for details).

In a representative DMN contact, ripple events elicited a clear burst of HFB activity (**Fig. 2A**). Hippocampal ripples induced a significant increase in HFB power in both AUT and OCT. Notably, DMN contacts showed greater ripple-aligned HFB power in the AUT than in the OCT (β = 0.008 ± 0.001, *P* < 10^-8^; **fig. S1**). This difference was specific to the ripple time window: when HFB power was averaged over the entire task period without regard to ripples, no significant difference between the AUT and OCT was observed (β = 0.003 ± 0.004, *P* = 0.506; **fig. S1**). Among DMN subregions (**Fig. 2B**), the ripple-aligned HFB increase in the AUT (relative to OCT) was most pronounced in the mPFC (β = 0.011 ± 0.002, *P*_FWE_ < 10^-4^; **Fig. 2C**) and was also evident in the lateral temporal cortex (β = 0.009 ± 0.002, *P*_FWE_ < 0.001). This pattern likely reflects the greater engagement of semantic schema-based processing required by the AUT (Beaty & Kenett, 2023; Ding et al., 2024; He et al., 2021; Kizilirmak et al., 2019), in contrast to the more feature-based processing of the OCT.

**Fig. 2.**
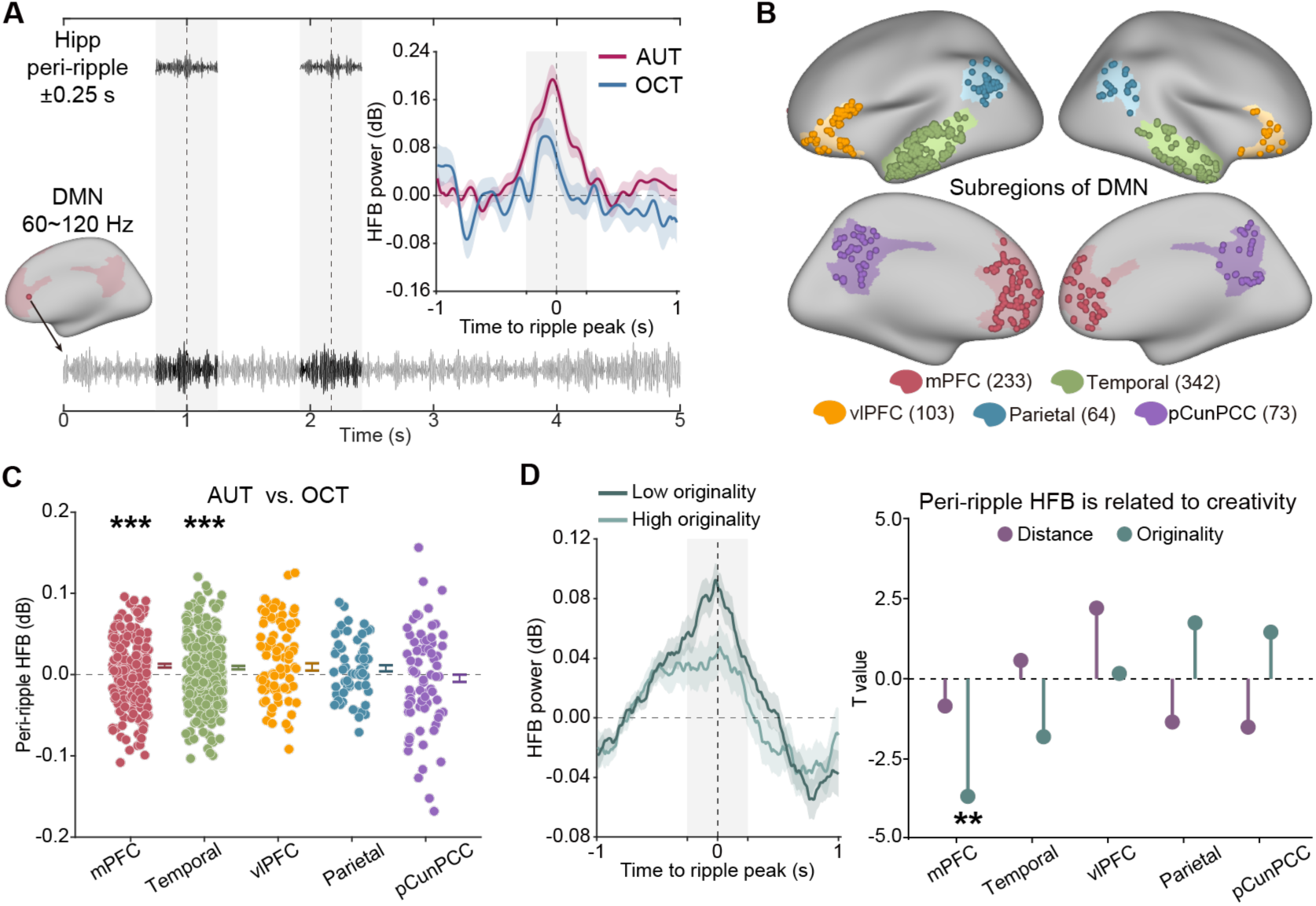
Ripple-aligned high-frequency activity in the mPFC supports creative thinking. (**A**) High-frequency broadband (HFB, 60–120 Hz) activity recorded from a DMN electrode (Subject 15, channel A’1) showed an increase of activity, time-locked to hippocampal ripple peaks during an example 5-second window (886.08–891.08 s after task onset). Inset: Ripple-aligned HFB power from a DMN electrode in the AUT (red) and OCT (blue); the grey shaded area denotes the ±250 ms peri-ripple window. (**B**) Anatomical definition of DMN subregions. Left: intracranial recording contacts were classified into DMN subdivisions using a 7-network atlas (Schaefer et al., 2018), including mPFC, ventrolateral PFC (vlPFC), lateral temporal cortex, parietal cortex, and precuneus/posterior cingulate (pCun/PCC). (**C**) Among DMN subregions, the mPFC and lateral temporal cortex showed significantly higher ripple-aligned HFB power in the AUT compared to the OCT. Data show mean (AUT vs. OCT) ± s.e.m, estimated using an LME model. Dots represent individual contacts for illustration purposes. (**D**) Left: Ripple-aligned HFB power from an mPFC electrode (Subject 38, channel K4) for AUT responses with high versus low originality. Right: Among DMN subregions, lower ripple-aligned HFB power in the mPFC was associated with higher AUT response originality. \*\**P* < 0.01; \*\*\**P* < 0.001.

Crucially, we found that moments of higher creativity were associated with a relative suppression of mPFC activity at the time of hippocampal ripples. Although ripple events in the AUT elicited overall higher mPFC HFB responses than those in the OCT (this was true even when considering high-originality and low-originality trials separately; both *P* < 0.001), within the AUT condition the most original ideas occurred when the ripple-aligned mPFC activation was relatively weak. In other words, higher response originality was predicted by lower ripple-aligned mPFC HFB power (β = - 0.006 ± 0.002, *P*_FWE_ = 0.001; **Fig. 2D**). This dissociation suggests that while the AUT broadly engages schema-related mPFC processes, excessive mPFC activation at the moment of a ripple may be counterproductive for generating original ideas.

Intriguingly, all of the above effects converged within the mPFC, revealing a notable anatomical specificity. Given the functional heterogeneity of the mPFC and the well-established link between the ventromedial PFC (vmPFC) and schematic representations (Ciaramelli et al., 2019; Gilboa & Marlatte, 2017; Klein-Flügge et al., 2022), we subdivided the mPFC contacts into vmPFC and dorsomedial PFC (dmPFC) regions using the Brainnetome atlas(Fan et al., 2016) (**fig. S2A**). The increase in ripple-aligned HFB for AUT vs OCT was significant in the vmPFC (β = 0.009 ± 0.004, *P* = 0.033; **fig. S2B**), but not in the dmPFC (β = 0.010 ± 0.005, *P* = 0.068). Likewise, the negative correlation between ripple-aligned mPFC activity and response originality was present only in the vmPFC (β = –0.007 ± 0.003, *P* = 0.010; **fig. S2C**) and was absent in the dmPFC (β = –0.005 ± 0.004, *P* = 0.137).

### Creative responses are linked to divergent mPFC representations during ripples

We next asked whether neural representations in the mPFC were reorganised during the formation of creative ideas – as would be expected if hippocampal ripples help liberate the mind from dominant schemas. Using a representational similarity analysis (RSA), we constructed multivariate activity patterns for high- and low-creativity responses during both the entire thinking period and the ripple-aligned window for a given object, pooling mPFC HFB signals across multiple contacts and participants. We then assessed the similarity of each condition-specific pattern to the overall task pattern, and compared these similarities between high- and low-creativity responses. The overall task pattern was defined as the mean HFB pattern across the entire AUT period for a given object (e.g., the average activity for “watermelon”), serving as a proxy for the schema-like representation of that task context (**Fig. 3A**; see Methods for details).

**Fig. 3.**
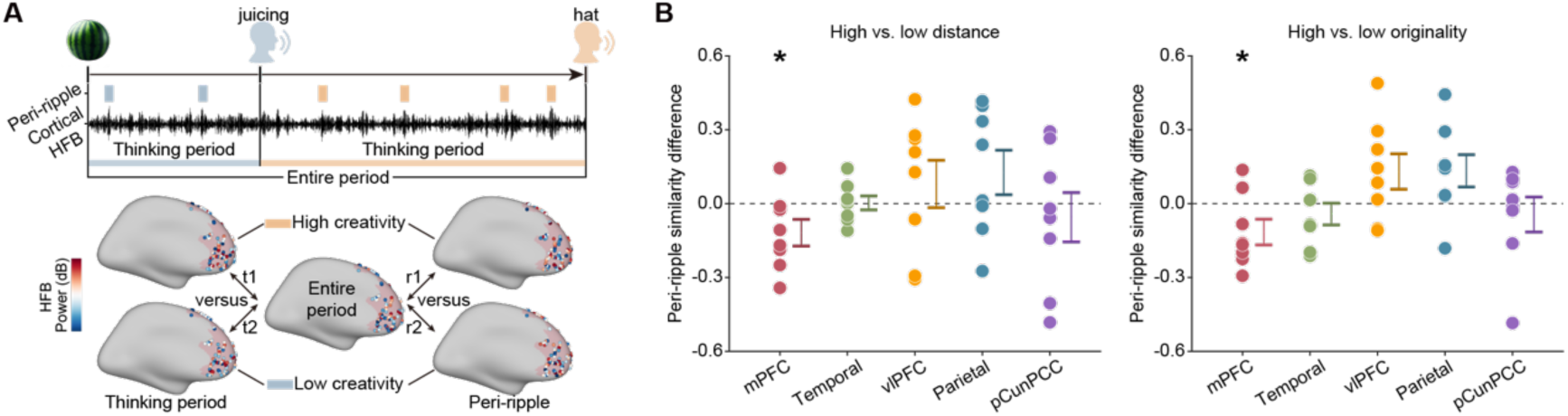
Ripple-aligned representational divergence in the mPFC support creative thinking. (**A**) Schematic of the representational similarity analysis. For each AUT object (e.g., watermelon), we compared neural activity patterns for high- vs low-creativity responses (defined by semantic distance or originality) during ripple events (“r1” vs “r2”) and during the thinking period (“t1” vs “t2”), relative to the overall task-average pattern. HFB power from all contacts in a given region (pooled across participants) was used to compute each pattern. For example, “r1” denotes the Pearson correlation between the region’s overall task pattern and its pattern for high-creativity responses during the peri-ripple window. “t1” denotes the Pearson correlation between the region’s overall task pattern and its pattern for high-creativity responses during the thinking period. (**B**) Among DMN subregions, only the mPFC showed a reduction in similarity between ripple-period neural representations and the overall task pattern for high- vs low-creativity responses. For illustration purposes, data show mean difference (high vs. low creativity) ± s.e.m, dots represent individual AUT objects. \**P* < 0.05.

During ripple-aligned windows, neural patterns in the mPFC for highly creative responses diverged significantly more from the overall task pattern than did those for less creative responses (semantic distance: *P* = 0.027; originality: *P* = 0.027; Wilcoxon test; **Fig. 3B**). This effect was evident for 7 of the 8 test objects in the AUT task; specifically, the neural patterns for highly creative ideas had consistently lower correlations with the task average than less creative ideas did (**Fig. 3B**). In contrast, no such divergence occurred when averaging over the entire thinking period without regard to ripple timing (semantic distance: *P* = 0.371; originality: *P* = 0.273).

To determine whether this peri-ripple representational shift was specific to creative ideation, we performed the same analysis on the control OCT data: no such ripple-related difference was observed in the OCT (semantic distance: *P* = 0.844). In other words, hippocampal ripples transiently reconfigured mPFC activity away from the dominant task representation when the ensuing idea was highly creative, whereas no such change occurred in the absence of a ripple or in non-creative contexts. This finding indicates that hippocampal ripple can briefly reshape mPFC representations away from existing schemas, facilitating the generation of original, non-standard ideas.

Further localising this effect within the mPFC, we separately examined the vmPFC and dmPFC. The greater representational divergence (high- vs low-creativity) during the ripple-aligned window was driven by the vmPFC (semantic distance: *P* = 0.039; originality: *P* = 0.055; **fig. S2D**), with no corresponding effect in the dmPFC (semantic distance: *P* = 0.273; originality: *P* = 0.156). These findings indicate that the vmPFC – a region strongly associated with schematic knowledge, is the key in which hippocampal ripples exert their creativity-modulating influence.

### Ripple-aligned mPFC activity tracks semantic switching along the dorsal–ventral axis

Such transient disengagement from schematic representations may also be necessary for flexibly traversing conceptual space during creative thought (Beaty & Kenett, 2023; Mastria et al., 2021; Nour et al., 2023). To test this possibility, we analysed how hippocampal ripples and ripple-aligned cortical activity related to semantic category transitions between consecutive ideas.

Within each AUT trial, we classified each response as either a category “clustering” event (staying within the same semantic category as the previous idea) or a category “switching” event (shifting to a new category), based on independently defined use categories for each object (see **Fig. 4A** and Methods). These classifications were verified by trained raters (e.g., for the object “umbrella”, using it for “shelter from rain” followed by “shelter from wind” constitutes clustering, whereas “shelter from wind” followed by “walking stick” represents a switch; see Methods for details). As expected, category switching was associated with greater creativity: switch responses showed higher semantic distance, higher originality ratings, and elevated hippocampal ripple rates compared to clustering responses (all *P* < 0.001; **Fig. 4B**).

**Fig. 4.**
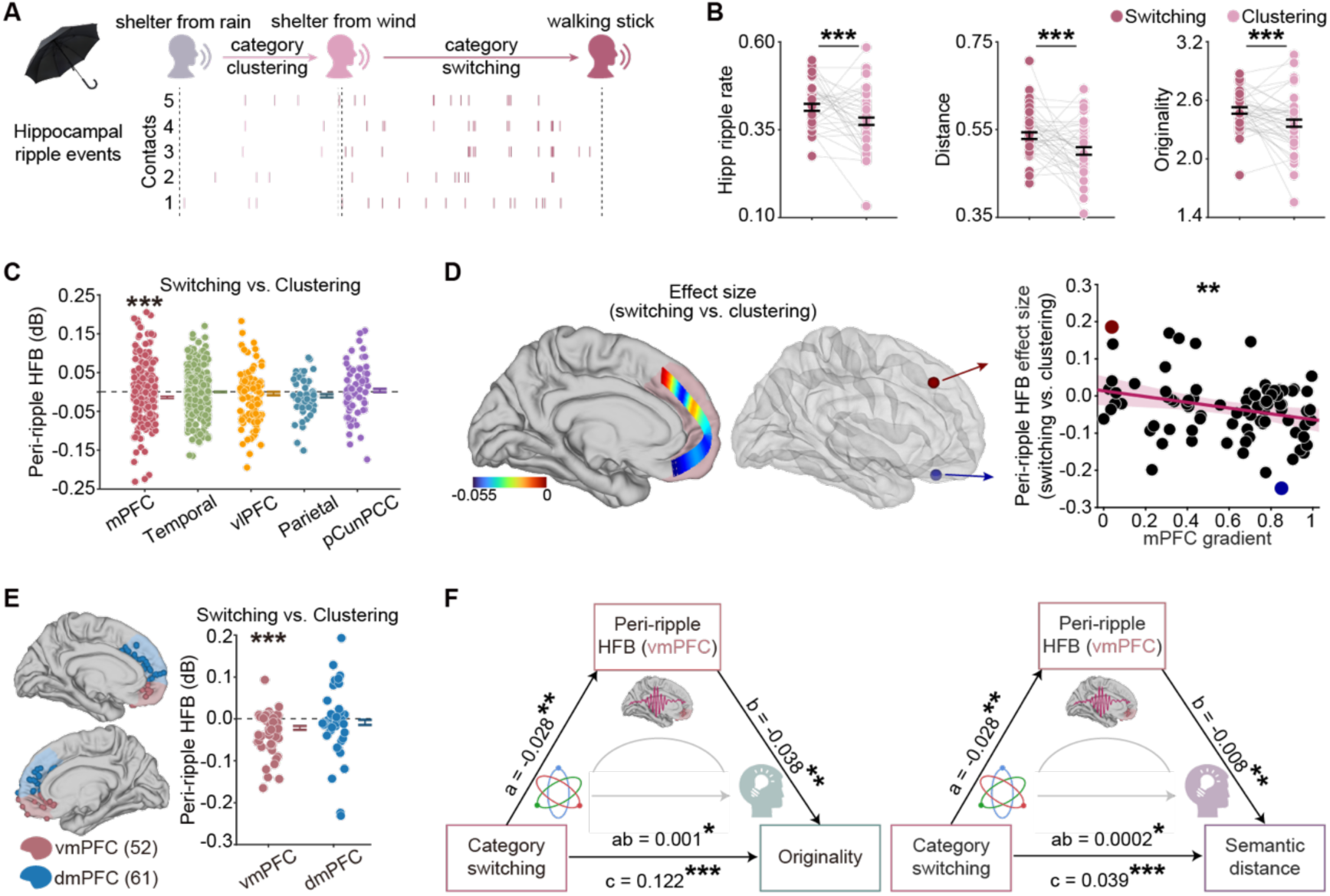
Ripple-aligned mPFC activity supports category switching during creative thinking (dorsal–ventral gradient). (**A**) Illustration of category clustering vs switching in the AUT (Subject 36, object: umbrella). Here, “shelter from rain” → “shelter from wind” represents a clustering transition (same functional category of use), whereas “shelter from wind” → “walking stick” is a switching transition (different category). Tick marks denote times of detected hippocampal ripples on five hippocampal contacts. (**B**) Compared to clustering, switching responses were accompanied by a higher hippocampal ripple rate, greater semantic distance, and higher originality scores. Data show mean ± s.e.m, estimated using an LME model. Dots represent individual subjects for illustration purposes. (**C**) Compared to clustering, switching responses showed lower ripple-aligned HFB power in the mPFC. Data show mean difference (switching vs. clustering) ± s.e.m, estimated using an LME model. Dots represent individual contacts for illustration purposes. (**D**) Left: Map of ripple-aligned HFB effect sizes for switching vs clustering across all mPFC contacts, plotted on a dorsal (top) to ventral (bottom) gradient. Warmer colours (red) indicate contacts where HFB was higher for clustering, cooler colours (blue) indicate higher HFB for switching. Middle: Two example mPFC contacts (large, coloured dots, one dorsal, one ventral) illustrate that ventral contacts exhibit a greater suppression of HFB during switching than dorsal contacts. Right: Across mPFC contacts, the switching–clustering HFB effect size was negatively correlated with the contact’s dorsal–ventral position, indicating the strongest switching-related suppression at the ventral end. Shaded areas indicate the 95% confidence interval of the fitted regression line. Dots represent individual mPFC contacts. (**E**) The vmPFC showed a significantly larger drop in ripple-aligned HFB power during switching compared to clustering, whereas the dmPFC did not. Data show mean difference (switching vs. clustering) ± s.e.m, estimated using an LME model. Dots represent individual contacts for illustration purposes. (**F**) Mediation analysis showing that the effect of category switching on response creativity was mediated by ripple-aligned vmPFC activity. The path from switching to higher creativity was significantly indirect via reduced ripple-aligned vmPFC activity. No significant mediation was found for the dmPFC (**fig. S5**). \**P* < 0.05, ** *P* < 0.01, *** *P* < 0.001.

Category switching was specifically accompanied by a suppression of mPFC activity time-locked to hippocampal ripples. During switching (vs clustering) events, ripple-aligned HFB power in the mPFC was markedly lower (β = −0.014 ± 0.003, *P*_FWE_ < 10^-7^; **Fig. 4C**). In line with our earlier results, this effect followed a clear dorsal–ventral gradient: the reduction in ripple-aligned HFB power with switching was most pronounced at more ventral mPFC contacts. Quantitatively, the magnitude of the switching-induced HFB power decreases at each mPFC contact correlated with that contact’s position along the dorsal–ventral axis (β = −0.077 ± 0.028, *P* = 0.007; **Fig. 4D**; see also **fig. S3**), indicating that the strongest ripple-linked suppression during idea switching occurred in the ventral mPFC. Notably, when we examined HFB power averaged over the entire thinking period (i.e., without aligning to ripples), none of these differences were present — there were no significant switching vs clustering differences in any DMN subregion or along the mPFC gradient (**fig. S4**). This confirms that the facilitation of idea switching is specifically tied to ripple-aligned cortical dynamics, rather than reflecting a general difference in sustained neural activity.

To further probe this effect, we separately analysed ripple-triggered responses in the vmPFC and dmPFC. The vmPFC showed a highly significant decrease in peri-ripple HFB power for category switching compared to clustering (β = −0.022 ± 0.005, *P* < 10^-6^; **Fig. 4E**), whereas the dmPFC showed no such effect (β = −0.008 ± 0.006, *P* = 0.187).

This vmPFC-specific suppression during switching prompted us to ask whether the ripple-aligned vmPFC activity might be the link between category switching and enhanced creativity. We performed a mediation analysis with the presence of category switching as the independent variable, creative performance (originality or semantic distance) as the outcome, and peri-ripple HFB power in the vmPFC as the mediator, controlling for overall hippocampal ripple rate and overall vmPFC HFB power. Indeed, this analysis revealed a significant mediation effect: part of the influence of category switching on creative performance was carried by the degree of ripple-aligned vmPFC suppression. Specifically, for originality score the mediated (indirect) effect was significant (z = 2.08, *P* = 0.037, 95% CI = [0.0002, 0.0020]), and the same was true for semantic distance (z = 1.98, *P* = 0.047, 95% CI = [4.45 × 10⁻⁵, 0.0005]; **Fig. 4F**). In contrast, the parallel mediation analysis for the dmPFC showed no significant effect (originality score: z = 0.43, *P* = 0.664, 95% CI = [-0.0005, 0.0010]; semantic distance: z = 0.46, *P* = 0.644, 95% CI = [-0.0001, 0.0003]; see also **fig. S5**). These results implicate a specific pathway: hippocampal ripples facilitate shifts in conceptual space (i.e., category switching) by transiently disengaging vmPFC activity, which in turn allows more creative, far-reaching ideas to emerge.

## Discussion

By leveraging simultaneous iEEG recordings from the hippocampus and multiple cortical regions, this study demonstrates that hippocampal ripples orchestrate prefrontal dynamics to foster creative idea generation. We found that coming up with more creative ideas was marked by an increase in hippocampal ripple events, a relative decrease in mPFC high-frequency activity coinciding with those ripples, and a divergence of ripple-time mPFC activity patterns away from the task’s typical (schema-like) state. Furthermore, this momentary reduction in ripple-aligned mPFC engagement facilitated dynamic switching between semantic categories of ideas, promoting exploration of novel associations. Notably, all of these effects followed a dorsal–ventral gradient within the mPFC, with the most pronounced changes in the vmPFC.

Hippocampal ripples are traditionally viewed as electrophysiological signatures of memory consolidation and planning (Buzsáki, 2015). Our results extend this framework by highlighting a role for ripples in creative thought. In rodents, hippocampal ripples coordinate the neural replay of previously experienced trajectories, illustrating its capacity to flexibly recombine stored elements depending on current task demands (Jadhav et al., 2012; Pfeiffer & Foster, 2013). Analogously, during human creative thinking, ripples may enable the recombination of remote semantic or conceptual traces, helping the mind leap from conventional ideas to novel solutions (Aru et al., 2023). Moreover, the observed correlation between ripple rates and individuals’ everyday creative activities suggests that this hippocampal mechanism is not confined to controlled laboratory settings, but may also underlie real-world creative expression across diverse domains.

Our findings on ripple-aligned mPFC activity help bridge a gap between the neuroscience of memory schemas and the neuroscience of creativity (Becker & Cabeza, 2025; Lewis et al., 2018). Past research has shown that hippocampal–prefrontal interactions can promote efficient memory use and schema-congruent inference (Preston & Eichenbaum, 2013; Sekeres et al., 2024; Xiao et al., 2025). However, our data indicate that in the context of creativity, a different dynamic may be beneficial: brief decoupling of the hippocampus–mPFC circuit. Indeed, rodent studies have demonstrated that sustained hippocampal–mPFC coupling can bias animals toward habitual behaviours, whereas introducing novelty to disrupt that coupling allows a switch to more flexible strategies (Park et al., 2021). This parallel suggests that transiently disengaging the hippocampal–prefrontal circuit can help break the mind out of rigid thought patterns. Our study advances this understanding by identifying hippocampal ripples as brief but pivotal events that modulate mPFC activity during creative cognition.

We propose that hippocampal ripples facilitate creativity by temporarily loosening the mPFC schema-based constraints (without turning them off completely). In doing so, ripples may allow access to ideas that would typically be suppressed under strong top-down schematic control. This mechanism aligns with classic theories in psychology which argue that relaxing the constraints of conventional knowledge is necessary for creativity to flourish (Duncker, 1945; Knoblich et al., 1999). Consistent with this, we observed that ripple events were accompanied by shifts in mPFC representations, supporting the notion that the brain can flexibly remap prefrontal activity patterns when exploring “out-of-the-box” ideas (Becker et al., 2025; H. Song et al., 2025).

Our dorsal–ventral gradient analysis highlights that the ripple-related effects were largely confined to the vmPFC. This anatomical specificity does not imply that the vmPFC operates alone; rather, it suggests that the vmPFC serves as a crucial focal point where hippocampal ripples influence schema-related processing during creative thinking. The vmPFC has been identified as a key hub for abstract, schematic knowledge (Gilboa & Marlatte, 2017; Sekeres et al., 2024), potentially providing a cortical substrate through which ripple events trigger transient changes in conceptual representations (H. Song et al., 2025). By moving beyond a coarse characterisation of “mPFC involvement,” our study, utilising iEEG, offers a more spatially refined account of how hippocampal–prefrontal dynamics support creative thought.

Finally, our findings open several avenues for future research. One practical direction is the development of closed-loop neurostimulation strategies to enhance creative thinking. For instance, a system could monitor hippocampal activity in real time and deliver a brief inhibitory stimulation to the vmPFC whenever a ripple is detected, to test whether this intervention boosts the originality of ideas. Additionally, since hippocampal ripples often co-occur with neural replay of past experiences (Buzsáki, 2015; Vaz et al., 2020), it will be interesting to investigate whether ripple-associated replay events help recombine memory elements in novel ways (Aru et al., 2023; Hahamy et al., 2023; Lewis et al., 2018). Such studies would further elucidate the hippocampal mechanisms that contribute to the human creative process.

## Methods

### Subjects

Intracranial recordings were acquired from 40 subjects (29 males and 11 females; age range: 11–54 years; mean age: 25.48 years; see **Table S1** for individual details) with refractory epilepsy who had depth electrodes implants for neurosurgical treatment at Sanbo Brain Hospital of Capital Medical University (Beijing, China). No clinical seizures occurred during the experiment. All participants provided informed consent. The study was approved by the institutional ethics review board (SBNK-YJ-2023-002-01) and conducted in accordance with the Declaration of Helsinki.

### Task and behavioural measures

Subjects completed a classic creative thinking paradigm, the Alternative Uses Task (AUT) (Guilford, 1967; Torrance, 1962), and a control task called the Object Characteristics Task (OCT) (Fink et al., 2009). In the AUT, subjects were given a common everyday object (e.g., watermelon or umbrella) and asked to generate as many unusual, creative, but also sensible uses for it as possible within 2 minutes. In the OCT, subjects were given a common object (e.g., toothbrush or cup) and asked to list as many typical features or characteristics of the object as possible within 1 minute. Eight AUT and eight OCT trials were administered per subject in a randomised order, following a few practice trials for task familiarisation. During each trial, subjects were instructed to remain silent while thinking and then to vocalise each idea immediately when it came to mind. All verbal responses were recorded via microphone. After the experiment, every response was transcribed and time-stamped (onset and offset) using the audio recording’s amplitude envelope, with manual checks to ensure accuracy. These timestamps were then used to align responses with the neural data for subsequent analyses.

Each AUT response was evaluated for originality by three trained raters using a standardised scoring guide (Beaty et al., 2018). The raters first agreed on whether the response was valid (i.e., a sensible use of the object). All responses deemed valid by unanimous agreement were then rated on a 5-point originality scale (1 = not original, 5 = highly original). The three ratings were averaged to yield the final originality score for each response. Inter-rater reliability was good (intraclass correlation coefficient, ICC= 0.68). Examples of responses with high and low originality scores are provided in **Table S2**. As an objective complement to human ratings, we also computed the semantic distance of each AUT response from its object prompt. Semantic distance was defined as 1 minus the cosine similarity between the word embedding of the response and that of the target object. We obtained 100-dimensional word embeddings from a large-scale Chinese corpus (Tencent AI Lab) (Y. Song et al., 2018). This metric captures how “far” a given idea is from the object typical associations. We found that semantic distance correlates positively with human-rated originality, indicating that it is a reasonable proxy for novelty.

To characterise idea transitions during the AUT, all responses for each object were grouped into functional use categories by three trained raters in collaborative discussion. We deliberately avoided clustering algorithms based on semantic distance, since embedding proximity does not always reflect functional similarity in high-dimensional conceptual space. For example, “doorstop” and “paperweight” are both ways of using a brick to hold something in place—functionally similar—but may lie far apart in embedding space and so would not cluster together. Such cases underscore the advantage of human judgment in identifying true categorical similarities. Using these curated categories, we then determined for each consecutive pair of responses whether they remained in the same category (category clustering) or shifted to a new one (category switching). For instance, with “umbrella,” moving from “shelter from rain” to “shelter from wind” was classified as clustering (both uses provide shelter), whereas transitioning from “shelter from wind” to “walking stick” was classified as switching (i.e., a new functional use).

In addition, subjects completed two questionnaires to assess everyday creativity and general fluid intelligence. Everyday creativity was measured using the frequency of everyday creative activities questionnaire, which evaluated the engagement in creative activities across nine domains (e.g., literature, music, handicrafts) over the past 12 months (Benedek et al., 2020). For each domain, subjects rated how often they engaged in creative activities during their leisure time on a 5-point scale: 0 = practically never, 1 = sometimes (a few times per year), 2 = regularly (about once a month), 3 = frequently (about once a week), and 4 = very frequently (nearly every day). Total scores provided an index of everyday creative activities. Fluid intelligence was assessed using Raven’s Standard Progressive Matrices (Raven, 2000). Participants were given 40 minutes to solve 60 items. Each item consisted of a 3 × 3 matrix with one missing piece, and subjects were required to select the correct completion from eight alternatives. The total number of correct responses served as the measure of fluid intelligence.

### Intracranial EEG recordings

iEEG data were recorded using a NeuroPort system (Blackrock Microsystems, Salt Lake City, UT, USA). Signals were band-pass filtered from 0.3 Hz to 500 Hz during acquisition and sampled at 2000 Hz. Each depth electrode probe (Sinovation, Beijing) had a diameter of 0.8 mm and carried 8 to 16 recording contacts (2 mm contact length, 1.5 mm inter-contact spacing). All electrode implantation sites were determined exclusively based on clinical needs.

### Electrode anatomical localisation

For each subject, a high-resolution T1-weighted MRI scan (pre-implantation) was processed with FreeSurfer (v7.3.2) (Fischl, 2012), to reconstruct the cortical surfaces. A post-implantation CT scan was then co-registered to the MRI (using the Brainstorm toolbox (Tadel et al., 2011) and SPM12 (Friston et al., 2006)) to visualise electrode locations. Each electrode contact was identified and manually marked on the co-registered images. An experienced neurologist verified all contact locations; any contact found to lie more than 3 mm outside the cortical surface was excluded from analysis. For group-level visualisation, all contact coordinates were projected into a common space (either FreeSurfer’s “fsaverage” brain or MNI standard space) using the iELVis toolbox (Groppe et al., 2017).

Anatomical labels were assigned to each contact in the subject’s native brain space by referencing two atlases: a 7-network cortical parcellation (Schaefer et al., 2018) and the Brainnetome atlas (Fan et al., 2016). The DMN was defined based on the 7-network parcellation and included contacts within the mPFC, ventrolateral PFC, lateral temporal cortex, parietal cortex, and precuneus/posterior cingulate cortex (pCun/PCC). For finer analysis of the mPFC along its dorsal–ventral axis, we subdivided mPFC contacts according to the Brainnetome atlas (Fan et al., 2016): ventromedial PFC (vmPFC, comprising areas A11m and A14m) versus dorsomedial PFC (dmPFC, areas A9m and A10m). This finer parcellation allowed us to examine functional gradients along the dorsal–ventral axis of mPFC.

### Intracranial EEG Pre-processing

All iEEG signals were downsampled offline to 500 Hz for processing. To remove mains electricity noise, we applied narrow notch filters at 50 Hz and its harmonics (100, 150, 200 Hz) using zero-phase (bidirectional) finite impulse response (FIR) filters with a 3 Hz stopband. We then performed an automated artifact screening to remove channels with excessive noise. An electrode contact was flagged for exclusion if any of the following were true: (i) its raw voltage or first temporal derivative exceeded the 95th percentile of all contacts, or (ii) its root-mean-square amplitude was more than 5 standard deviations above the mean of all contacts. Every contact flagged by these criteria was then visually inspected (in both time-series and spectrogram forms) to confirm whether it was artefactual; confirmed noisy contacts were removed from further analysis.

The signals from remaining contacts were re-referenced using a bipolar montage: each contact’s time series was subtracted from that of an immediately adjacent contact on the same electrode probe. For data from hippocampal contacts, we performed bipolar referencing by subtracting each hippocampal contact’s signal from a neighbouring contact located in the white matter on the same probe, as determined by MRI segmentation (Fischl, 2012). This bipolar re-referencing minimises the influence of volume-conducted signals and focuses the data on local intracranial activity.

### Hippocampal ripples detection

Following a previously published protocol (Norman et al., 2019, 2021), hippocampal ripples were detected on bipolar iEEG recordings from contacts located within 3 mm of the CA1, CA2/CA3, and subiculum subfields, where ripples are most reliably observed (Chrobak & Buzsáki, 1996; Oliva et al., 2016). Individual hippocampal subfields were delineated from pre-implantation structural MRI scans using the FreeSurfer parcellation algorithm (Iglesias et al., 2015).

The bipolar iEEG signals were bandpass filtered between 70–180 Hz using a zero-lag linear-phase Hamming-windowed FIR filter with a 5 Hz transition bandwidth. The analytic amplitude was then extracted using the Hilbert transform. To minimise bias from transient artefacts or extreme values, amplitude outliers were clipped at 4 standard deviations above the mean using a robust estimator based on the least median of squares. The resulting signal was squared and smoothed using a Kaiser-window FIR low-pass filter with a 40 Hz cutoff, and its mean and standard deviations over the entire task were used to define the ripple detection threshold. Candidate ripple events were identified from the original (non-clipped) squared signal if they exceeded 4 standard deviations above the threshold. Events were extended until the ripple-band power fell below 2 standard deviations. Events shorter than 20 ms or longer than 200 ms were excluded, and temporally adjacent events with peak-to-peak separation less than 30 ms were merged. Finally, ripple peaks were aligned to the nearest trough in the raw (non-rectified) signal corresponding to the peak of ripple-band power.

To control for potential contamination by global artefacts, a control detection procedure was applied to the common average signal across all contacts. Ripple events coinciding with peaks in this global signal were excluded to mitigate the influence of transient electrical or muscular artefacts that may manifest across multiple channels simultaneously (Fiederer et al., 2016).

To further avoid inclusion of pathological activity, ripple events occurring within 100 ms of interictal epileptiform discharges were excluded (Gelinas et al., 2016). Interictal epileptiform discharges were detected by filtering hippocampal signals between 25–60 Hz (zero-lag linear-phase Hamming-windowed FIR filter), followed by rectification, squaring, smoothing, and thresholding at 4 standard deviations above the baseline.

In addition, hippocampal ripples were distinguished from pathological high-frequency oscillations (HFOs) that may occur in epileptic regions (Bragin et al., 2010). Hippocampal HFOs with characteristics indicative of epileptiform activity were detected using a validated method (S. Liu & Parvizi, 2019), and any ripple events occurring within 100 ms of such HFOs were removed from further analysis.

Based on the identified hippocampal ripples, ripple rate was calculated as the number of ripple events divided by the duration of time (in seconds), excluding periods of verbalisation.

### Deconvolution of peri-ripple cortical HFB activity

To analyse cortical activity around the times of hippocampal ripples, we focused on high-frequency broadband power (HFB, 60–120 Hz) as an index of local cortical activation. We filtered the pre-processed cortical signals (bipolar derivations) in the 60– 120 Hz band (using a linear-phase FIR filter with 5 Hz transition band) and extracted the analytic amplitude via Hilbert transform. The HFB amplitude was then converted to decibel (dB) units by 10·log^10^ for normalisation.

Given multiple events of interest can overlap in time (e.g., a hippocampal ripple may occur shortly before or after a verbal response), we employed a linear deconvolution approach to separate ripple-locked cortical activity from other event-related activity. We used the Unfold toolbox (Ehinger & Dimigen, 2019), to perform regression-based deconvolution on the continuous HFB power time series (downsampled to 100 Hz for computational efficiency). The model treated the observed HFB signal at each cortical contact as the sum of contributions from three types of events: (i) hippocampal ripple peaks, (ii) speech onset (response onset), and (iii) speech offset (end of vocalisation). Each event type’s impulse response was modelled using a set of 40 cubic spline basis functions spanning from 1 s before to 1 s after the event. This time-expanded design matrix allowed us to estimate a distinct peri-event time course (set of regression beta weights) for each event type, even when events overlapped in time. In essence, the deconvolution fits a unique temporal response function for ripple events, while accounting for any concurrent response-related activity. The resulting beta weights corresponding to ripples represent the isolated cortical HFB response time-locked to hippocampal ripples.

From these deconvolved ripple response functions, we defined a peri-ripple window of interest as –250 ms to +250 ms around the ripple peak. This time window was chosen to capture the immediate ripple-aligned cortical dynamics while minimising contamination from unrelated neural events, consistent with previous human ripple studies (Norman et al., 2021; Xiao et al., 2025). For each cortical contact, we averaged the ripple-related HFB beta values across this ±250 ms window to yield a single measure of ripple-aligned HFB magnitude. We then used LME models to assess differences in this peri-ripple HFB magnitude between conditions and to examine correlations with creativity measures.

### Representational similarity analysis

We examined whether higher creativity was associated with more divergent neural representations in the mPFC. Specifically, we asked whether the pattern of mPFC activity preceding high-creativity responses was less similar to the overall task context pattern than that preceding low-creativity responses.

For each subject and each AUT object, responses were divided into “high creativity” and “low creativity” groups based on their originality scores or semantic distances (analysed separately). We then computed the mean HFB power for each contact within a given region across three time windows: (i) the thinking period prior to response onset, (ii) the peri-ripple window (±250 ms around the ripple peak), and (iii) the entire task duration excluding speech periods (used to defined the overall task pattern). Next, we pooled contacts across all subjects within each DMN subregion to construct region-level activity patterns.

We then quantified the similarity between each condition-specific pattern (from the thinking and peri-ripple periods) and the overall task pattern using Pearson correlation. Specifically, we obtained four correlation coefficients for each AUT object: *r1* = correlation between high-creativity pattern during the peri-ripple period and the overall task pattern; and *r2* = correlation between low-creativity pattern during the peri-ripple period and the overall task pattern; *t1* = correlation between high-creativity pattern during the thinking period and the overall task pattern; *t2* = correlation between low-creativity pattern during the thinking period and the overall task pattern (**Fig. 3A**). We focused on comparing the high- vs low-creativity similarities within each time window across objects (i.e., *r1* vs *r2*, *t1* vs *t2*).

### mPFC gradient analysis

We further analysed the mPFC results along a continuous dorsal–ventral anatomical gradient. We defined a curved axis along the medial wall of each hemisphere passing through the dmPFC and vmPFC regions (with gradient values normalized from 0 at the most dorsal point to 1 at the most ventral point). Each mPFC contact location was projected onto this axis to assign it a gradient value between 0 and 1, representing its dorsal–ventral position. We then quantified each contact’s switching vs clustering effect on ripple-aligned HFB by calculating the Cohen’s *d* effect size for the HFB difference between switching and clustering responses. We performed this calculation separately for the peri-ripple window and, for comparison, for the thinking period. Finally, we used robust linear regression to test for a relationship between the contact’s effect size and its anatomical gradient position. A significant negative correlation would indicate that the effect (switching vs clustering) is weaker at one end of the mPFC (ventral) and stronger at the other (dorsal), consistent with the observations from the vmPFC vs dmPFC categorical analysis.

### Mediation analysis

We conducted a mediation analysis focusing on ripple-aligned mPFC activity as a mediator. In this analysis, the independent variable (X) was the extent of category switching on a trial (a binary indicator for a switching event when analyzing individual transitions), and the dependent variable (Y) was the creative performance outcome (either the originality rating or semantic distance of response). The mediator (M) was the peri-ripple HFB power in the vmPFC or dmPFC during the response’s thinking period. We included covariates for the overall hippocampal ripple rate and the overall mean cortical HFB power (to ensure that the mediation specifically involved the ripple-aligned activity, beyond any general differences in ripple frequency or baseline activation).

We followed standard mediation analysis procedures: we computed the total effect of X on Y (path c), the effect of X on M (path a), and the effect of M on Y while controlling for X (path b). The indirect (mediated) effect was quantified as a × b. We used a Monte Carlo resampling approach (1,000 iterations, quasi-Bayesian) to estimate 95% confidence intervals for the indirect effect as well as for the other path coefficients. Mediation was considered significant if (i) the confidence intervals for both the total and indirect effects excluded zero, and (ii) the total and indirect effects had the same sign, indicating a consistent direction of influence via the mediator. The mediation analysis was implemented in R v4.3.1 using the bruceR package (Bao, 2024).

### Statistical analysis

We tested the effect of task (AUT vs OCT) on hippocampal ripple rate and peri-ripple HFB power, and similarly tested the effect of category (switching vs clustering). Given the nested data structure, with multiple hippocampal contacts recorded for each subject, LME models were used to account for variability across both subjects and contacts (Yu et al., 2022). When testing effects across multiple DMN subregions, Bonferroni correction was applied to control for multiple comparisons. These models were formulated as follows:

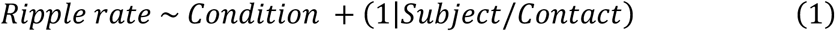

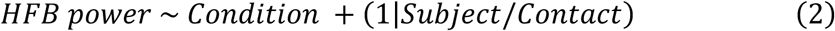

We also used LME models to assess whether hippocampal ripple rate and peri-ripple HFB predicted creative performance, indexed by originality scores and semantic distances. Notably, the predictive analysis for hippocampal ripple rate was conducted at the trial level rather than the response level, due to the sparsity of ripple events and the brevity of some thinking periods. Such short windows could yield either zero or disproportionately high ripple rates, thereby compromising the reliability of the estimates.

For representational similarity analysis, given our a priori hypothesis focusing on the mPFC, similarity differences were assessed using one-tailed Wilcoxon signed-rank tests across AUT objects, testing whether high-creativity responses were associated with lower pattern similarity (i.e., greater divergence from the overall task pattern) than low-creativity responses. Analyses in other DMN subregions were included for exploratory purposes and were not subjected to multiple comparison correction.

## Acknowledgments

We thank the patients for their participation and invaluable contribution to this research.

## Author contributions

Conceptualization: Y.L., L.H. Investigation: L.H., X.W., J.Z., Z.X., X.H., Y.L. Writing – original draft: L.H., Y.L. Writing – review & editing: L.H., Y.L.

## Data and materials availability

The statistical data and analysis code supporting the conclusions of this study will be made available on https://doi.org/10.5281/zenodo.15619058 upon publication.

## Supplementary Information

**Fig. S1.**
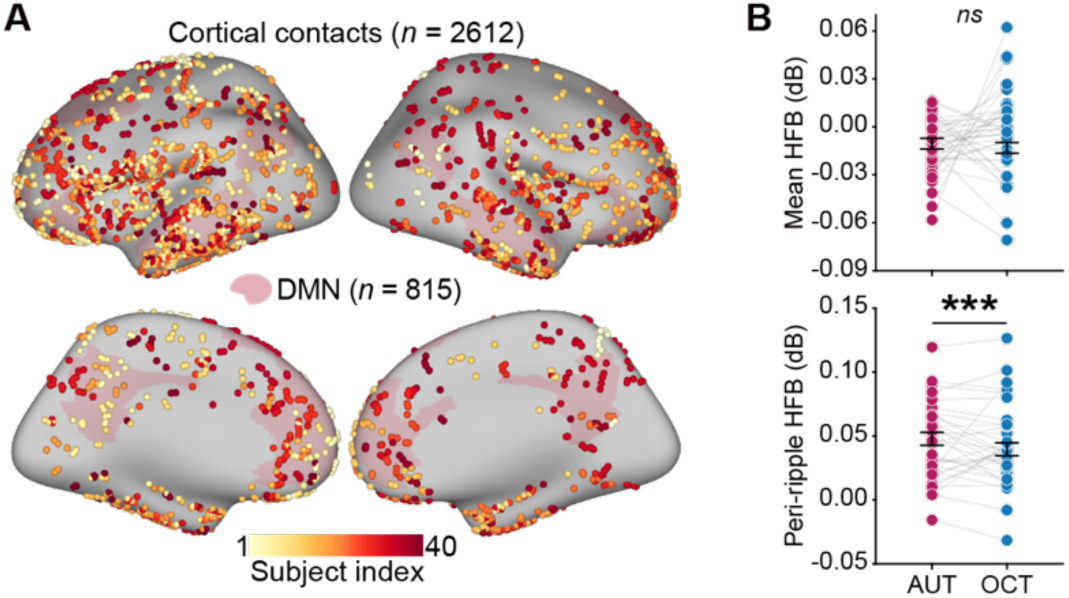
Ripple-aligned DMN activity during creative thinking. (**A**) Distribution of 2,612 cortical contacts across 40 subjects, including 815 contacts within the DMN. Each contact is colour-coded based on the subject index. (**B**) Peri-ripple HFB power was significantly greater during the AUT than the OCT (β = 0.008 ± 0.001, *P* < 10-8); however, when HFB was averaged across the entire task period without ripple alignment, no significant task-related differences were observed. Data show mean ± s.e.m, estimated using an LME model. Dots represent individual subjects for illustration purposes. \*\*\**P* < 0.001.

**Fig. S2.**
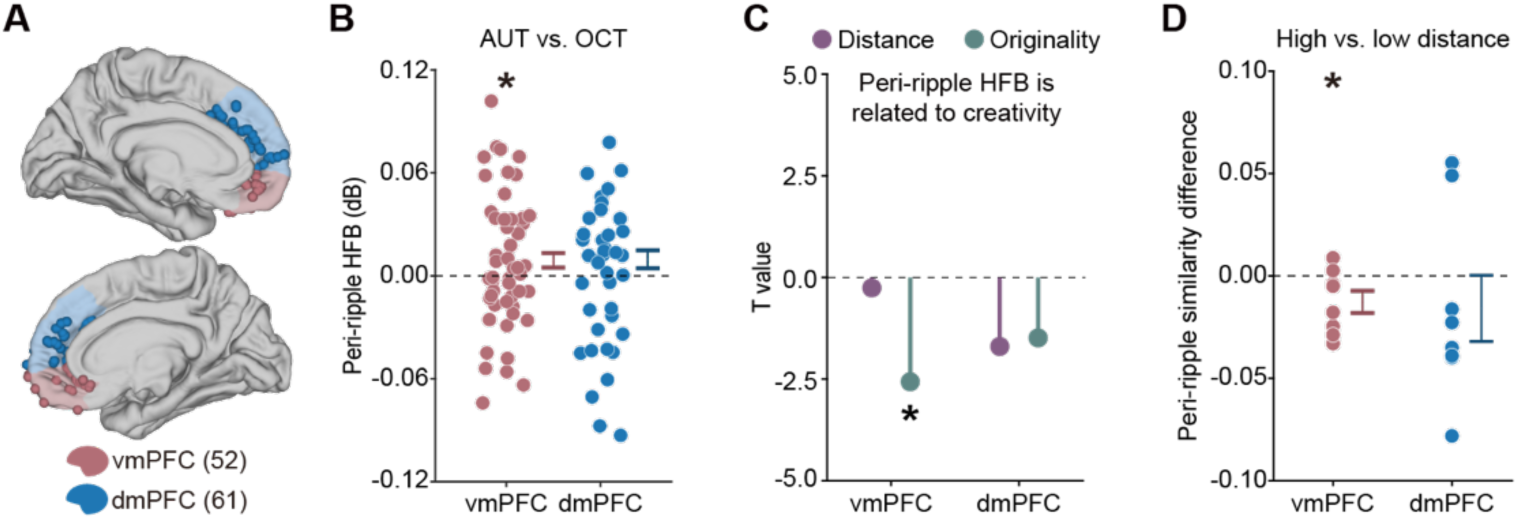
Hippocampal ripple-linked creative effects are localised to vmPFC. (**A**) The mPFC contacts were further subdivided into ventromedial (vmPFC) and dorsomedial (dmPFC) based on the Brainnetome atlas (*52*), delineating a clear dorsal–ventral axis. (**B**) The vmPFC showed significantly higher ripple-aligned HFB power in the AUT compared to the OCT. No significant difference was observed in the dmPFC. Data show mean difference (AUT vs. OCT) ± s.e.m, estimated using an LME model. Dots represent individual contacts for illustration purposes. (**C**) Lower ripple-aligned HFB power in the vmPFC was associated with higher AUT response originality. No significant correlation was observed in the dmPFC. (**D**) The vmPFC showed a reduction in similarity between ripple-period neural representations and the overall task pattern for high- vs low-distance responses. This representational divergence effect was not observed in the dmPFC. For illustration purposes, data show mean difference (high vs. low creativity) ± s.e.m, dots represent individual AUT objects. \**P* < 0.05.

**Fig. S3.**
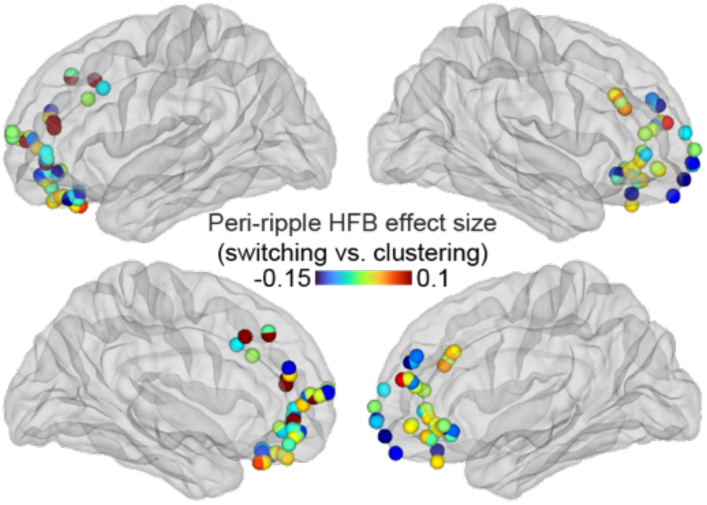
Ripple-aligned HFB effect sizes for switching versus clustering in mPFC contacts. Spheres mark individual mPFC contacts projected onto lateral (top row) and medial (bottom row) views of the left and right hemispheres. Colours encode the peri-ripple HFB effect size (switching vs. clustering), with warm hues (red) indicating greater HFB power during switching and cool hues (blue) indicating greater HFB power during clustering.

**Fig. S4.**
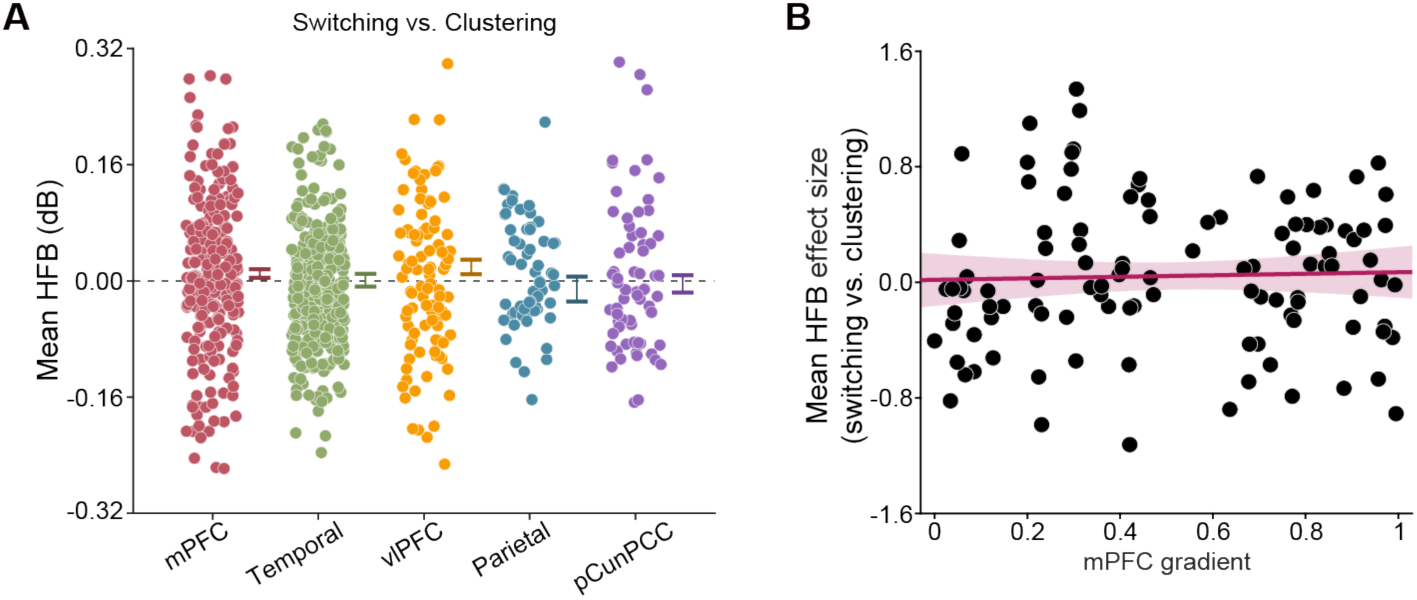
No significant switching versus clustering effects in mean HFB activity without ripple alignment. (**A**) Mean HFB power difference (switching vs. clustering) averaged over the thinking period without ripple alignment for five DMN subregions. No region exhibited a significant HFB difference (all *P*_FWE_ > 0.05). Data show mean difference (switching vs. clustering) ± s.e.m, estimated using an LME model. Dots represent individual contacts for illustration purposes. (**B**) Mean HFB effect sizes (switching vs. clustering) for individual mPFC contacts were not correlated with their anatomical gradient values along the dorsal–ventral axis (*P* > 0.05). Shaded areas indicate the 95% confidence interval of the fitted regression line. Dots represent individual mPFC contacts.

**Fig. S5.**
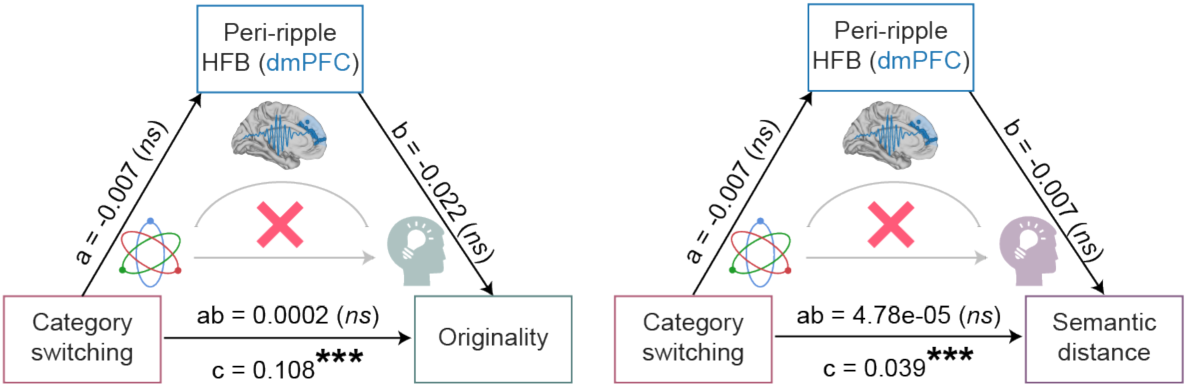
Lack of mediation by ripple-aligned dmPFC activity in the link between switching and creativity. Each panel depicts a non-significant mediation model for originality rating (mediated effect: z = 0.43, *P* = 0.66, 95% CI = [-0.0005, 0.0010]) and semantic distance (mediated effect: z = 0.46, *P* = 0.64, 95% CI = [-0.0001, 0.0003]), respectively.

**Table S1.** Demographic information and number of contacts for each subject.

| ID | Sex | Age | Handedness | Raven | Hipp | DMN | mPFC | vIPFC | Temporal | Partial | pCunPCC | vmPFC | dmPFC |
| --- | --- | --- | --- | --- | --- | --- | --- | --- | --- | --- | --- | --- | --- |
| 001 | Female | 13 | Right | N/A | 0 | 7 | 0 | 0 | 7 | 0 | 0 | 0 | 0 |
| 002 | Male | 18 | Right | N/A | 4 | 15 | 7 | 4 | 4 | 0 | 0 | 0 | 0 |
| 003 | Female | 19 | Right | 47 | 0 | 6 | 0 | 4 | 2 | 0 | 0 | 0 | 0 |
| 004 | Male | 15 | Right | 49 | 14 | 16 | 0 | 10 | 4 | 2 | 0 | 0 | 0 |
| 005 | Female | 18 | Right | 47 | 5 | 11 | 3 | 2 | 0 | 0 | 6 | 0 | 0 |
| 006 | Male | 11 | Right | 29 | 2 | 8 | 3 | 1 | 4 | 0 | 0 | 0 | 1 |
| 007 | Male | 29 | Right | 32 | 12 | 32 | 0 | 8 | 24 | 0 | 0 | 0 | 0 |
| 008 | Male | 35 | Right | 41 | 0 | 17 | 14 | 3 | 0 | 0 | 0 | 0 | 7 |
| 009 | Male | 17 | Right | 47 | 4 | 15 | 0 | 0 | 2 | 9 | 4 | 0 | 0 |
| 010 | Male | 25 | Right | 34 | 6 | 24 | 6 | 5 | 13 | 0 | 0 | 0 | 0 |
| 011 | Male | 23 | Right | 54 | 6 | 23 | 4 | 7 | 12 | 0 | 0 | 3 | 0 |
| 012 | Male | 22 | Right | 52 | 6 | 14 | 7 | 2 | 5 | 0 | 0 | 3 | 6 |
| 013 | Female | 14 | Right | 38 | 10 | 12 | 0 | 0 | 2 | 5 | 5 | 0 | 0 |
| 014 | Male | 15 | Right | 14 | 4 | 11 | 6 | 1 | 4 | 0 | 0 | 2 | 0 |
| 015 | Female | 33 | Left | 50 | 7 | 18 | 0 | 4 | 13 | 1 | 0 | 0 | 0 |
| 016 | Male | 49 | Right | 53 | 0 | 12 | 0 | 0 | 5 | 6 | 1 | 0 | 0 |
| 017 | Male | 30 | Right | 46 | 5 | 30 | 3 | 10 | 17 | 0 | 0 | 3 | 0 |
| 018 | Male | 30 | Right | 57 | 5 | 27 | 9 | 8 | 10 | 0 | 0 | 6 | 2 |
| 019 | Male | 26 | Right | 22 | 8 | 13 | 0 | 2 | 11 | 0 | 0 | 0 | 0 |
| 020 | Female | 42 | Right | 33 | 10 | 15 | 4 | 0 | 11 | 0 | 0 | 0 | 0 |
| 021 | Male | 18 | Right | 52 | 8 | 23 | 13 | 6 | 4 | 0 | 0 | 3 | 5 |
| 022 | Male | 19 | Right | 45 | 7 | 25 | 2 | 1 | 22 | 0 | 0 | 0 | 0 |
| 023 | Male | 34 | Right | 34 | 20 | 14 | 11 | 0 | 3 | 0 | 0 | 5 | 1 |
| 024 | Male | 12 | Ambidexterity | 8 | 0 | 9 | 5 | 4 | 0 | 0 | 0 | 0 | 2 |
| 025 | Male | 27 | Right | 24 | 0 | 23 | 16 | 1 | 6 | 0 | 0 | 0 | 13 |
| 026 | Female | 53 | Right | 41 | 5 | 24 | 10 | 1 | 8 | 3 | 2 | 4 | 6 |
| 027 | Male | 28 | Ambidexterity | 46 | 5 | 18 | 5 | 0 | 8 | 4 | 1 | 0 | 0 |
| 028 | Male | 31 | Right | 19 | 5 | 16 | 0 | 0 | 7 | 7 | 2 | 0 | 0 |
| 029 | Male | 18 | Right | N/A | 4 | 42 | 37 | 2 | 3 | 0 | 0 | 3 | 6 |
| 030 | Male | 19 | Right | 56 | 4 | 35 | 22 | 9 | 4 | 0 | 0 | 4 | 5 |
| 031 | Male | 30 | Right | 52 | 6 | 39 | 4 | 4 | 24 | 3 | 4 | 0 | 0 |
| 032 | Female | 14 | Right | 52 | 1 | 20 | 0 | 0 | 5 | 3 | 12 | 0 | 0 |
| 033 | Female | 26 | Left | 50 | 4 | 18 | 0 | 0 | 7 | 4 | 7 | 0 | 0 |
| 034 | Male | 16 | Right | N/A | 4 | 25 | 8 | 1 | 16 | 0 | 0 | 2 | 1 |
| 035 | Male | 26 | Right | 55 | 11 | 23 | 2 | 0 | 13 | 0 | 8 | 2 | 0 |
| 036 | Male | 27 | Right | 56 | 8 | 13 | 2 | 1 | 8 | 1 | 1 | 2 | 0 |
| 037 | Female | 54 | Right | 44 | 8 | 17 | 0 | 0 | 11 | 1 | 5 | 0 | 0 |
| 038 | Female | 16 | Right | 56 | 16 | 36 | 6 | 2 | 22 | 6 | 0 | 4 | 0 |
| 039 | Male | 37 | Right | 31 | 6 | 23 | 8 | 0 | 4 | 5 | 6 | 3 | 2 |
| 040 | Male | 30 | Right | 42 | 17 | 46 | 16 | 0 | 17 | 4 | 9 | 3 | 4 |

**Table S2.** Representative responses for high and low originality scores.

| Object | Five responses selected from the top 10% of originality scores | Five responses selected from the bottom 10% of originality scores |
| --- | --- | --- |
| Watermelon | Lamp<br>Haircut practice<br>Dumbbell<br>Helmet<br>Target | Buying and selling<br>Salad ingredient<br>Watermelon juice<br>Eating<br>Quenching thirst |
| Pop can | Lantern<br>Ring<br>Mailbox<br>Pocket knife<br>Stilts | Cup<br>Smashing<br>Holding beverages<br>Aluminium source<br>Recycling |
| Toothpick | Hair styling<br>Applying makeup<br>Puzzle piece<br>Arrow<br>Maze design | Cleaning<br>Picking teeth<br>Poking<br>Bamboo skewer<br>Piercing |
| Umbrella | Hovercraft<br>Speed brake<br>Plug the hole<br>Fan<br>Fishing tool | Buying and selling<br>Blocking light<br>Rain cover<br>Wind protection<br>Sunshade |
| Ruler | Lever<br>Slingshot<br>Ping pong paddle<br>Chopsticks<br>Fence | Craft material<br>Drawing lines<br>Drawing rectangles<br>Measuring length<br>Measuring height |
| Carton | Lantern<br>Pinwheel<br>Tank toy<br>Insulation container<br>Fan | Recycling<br>Folding<br>Storing goods<br>Carrying items<br>Lunchbox |
| Pillow | Simulating pregnancy<br>Swim ring<br>Hat<br>Emotional outlet<br>Protective gear | Cushion<br>Headrest<br>Sleeping<br>Lying down<br>Backrest |
| Hat | Begging prop<br>Building a snowman<br>Eye mask<br>Frisbee<br>Bird nest | Playing<br>Sunshade<br>Rain shield<br>Wind protection<br>Cold protection |

